# The maize *striate leaves2* (*sr2*) gene encodes a conserved DUF3732 domain homologous to the rice *yss1* gene

**DOI:** 10.1101/2023.10.05.561108

**Authors:** Meghan J. Brady, Maya Cheam, Jonathan I. Gent, R. Kelly Dawe

## Abstract

Maize s*triate leaves2* (*sr2*) is a mutant that causes white stripes on leaves that has been used in mapping studies for decades, though the underlying gene has not been identified. The *sr2* locus has been previously mapped to small regions of the normal chromosome 10 (N10) and a rearranged variant called Abnormal Chromosome 10 (Ab10). A comparison of assembled genomes carrying N10 and Ab10 revealed only five candidate *sr2* genes. Analysis of a stock carrying the *sr2* reference allele (*sr2-ref*) showed that one of the five genes has a transposon insertion that disrupts its protein sequence and has a severe reduction in mRNA. An independent Mutator transposon insertion in the gene (*sr2-Mu*) failed to complement the *sr2-ref* mutation, and plants homozygous for *sr2-Mu* showed white striped leaf margins. The *sr2* gene encodes a DUF3732 protein with strong homology to a rice gene with a similar mutant phenotype called *young seedling stripe1* (*yss1*). These and other published data suggest that *sr2* may have a function in plastid gene expression.

## INTRODUCTION

A large number of maize nuclear genes provide products necessary for chloroplast function. Mutations in these genes result in albino, virescent, pale green, yellow, or white striped plants. In the striped class there are at least eight different maize loci - *striate leaves1, striate leaves2, striate leaves3, striate leaves4, japonica striping1, japonica striping2, iojap striping1* and *iojap striping2* (Gerald Neuffer et al., 1997). These mutants impact the efficiency of plastid transcription, translation or morphogenesis such that chloroplast function is impaired but not abolished. Striping is likely caused by the sorting out of mixed populations of functional and non-functional chloroplasts in a way that some lineages inherit no functional chloroplasts and appear as white sectors (Birky, 1983; Coe et al., 1988). White stripes tend to be wider and more common at the margins of leaves, because cells at the margin undergo more division to expand the width of the leaf than cells in the center of the leaf (Walbot and Coe, 1979; Han et al., 1992; Park et al., 2000).

The *iojap striping1* (*ij1*) gene has been described at the molecular level (Rhoades, 1943; Han et al., 1992). Chloroplasts within white stripes of *ij1* mutants are present but lack ribosomes, suggesting a failure in ribosome assembly (Shumway and Weier, 1967; Walbot and Coe, 1979; Siemenroth et al., 1980). *Iojap* is a member of a conserved family of ribosomal silencing factor A/DUF143-containing proteins (Häuser et al., 2012; Wang et al., 2016) that function in bacterial and chloroplast ribosome biogenesis and protein synthesis (Walbot and Coe, 1979; Trösch and Willmund, 2019). More recently a mutant called *white and green striate leaves1 (wgsl1)* was described (Li et al., 2023), which may be an allele of *striate leaves4* on chromosome 6. The *wgsl1* gene encodes a 16S rRNA processing protein that is thought to be required for ribosome maturation (Li et al., 2023).

S*triate leaves2* (*sr2*) was originally identified in Waseca Minnesota as a spontaneous mutant (Joachim and Burnham, 1953). Electron microscope analysis of white tissue revealed chloroplasts with unorganized internal structure and a lack of visible ribosomes, similar to *ij1* (Williams and Kermicle, 1974). The *sr2* gene is located on the long arm of chromosome 10, both on the normal form of chromosome 10 (N10) and a variant of chromosome 10 known as abnormal chromosome 10 (Ab10) (Rhoades and Dempsey, 1985). Here we combine comparative genomics with genetic and molecular analyses of two alleles of the *sr2* gene to demonstrate that it encodes a DUF3732 protein with homology to rice *Young Seedling Stripe1 (Zhou et al*., *2017)*, a gene that is thought to modulate gene expression by plastid-encoded plastid RNA polymerase.

## RESULTS

### Comparative genomics of N10 and Ab10

We started our analysis with a careful comparison of two forms of chromosome 10, the normal chromosome 10 (N10) and abnormal chromosome 10 (Ab10). The two chromosomes are syntenic except at the ends of their long arms, where, on Ab10, there is a large ∼55 Mb meiotic drive haplotype (Liu et al., 2020; Dawe, 2022). The Ab10 meiotic drive haplotype causes the preferential transmission of Ab10 when crossed as a female (Dawe, 2022). Portions of the end of normal chromosome 10 (N10) are present within the Ab10 haplotype, though the order of genes is altered by inversions and rearrangements (Rhoades and Dempsey, 1985). On N10, four genes with visible mutant phenotypes called *white2* (*w2*), *opaque7* (*o7*), *luteus13* (*l13*) and *striate leaves2* (*sr2*) occur in the order *w2-o7-l13-sr2*, whereas on Ab10 the gene order is *l13-o7-w2-sr2* (Rhoades and Dempsey, 1985). More comprehensive mapping demonstrated that there are two separate inversions but that *sr2* is not included in either one (Mroczek et al., 2006). Complete genome assembly of the Ab10 haplotype (Liu et al., 2020) and subsequent whole genome alignments confirmed the two inversions but did not identify an obvious uninverted region of homology at the ends of the shared region (Liu et al., 2020; Song et al., 2022), raising concerns about the original interpretations of gene order.

To confirm the location of *sr2*, we grew and analyzed a line homozygous for the Ab10 terminal deletion line known as Ab10-Df(K). Ab10-Df(K) had been described as having the *sr2* phenotype and a reduced stature, but otherwise appearing normal, suggesting that only a small section of the Ab10 shared region, including *sr2*, was missing (Rhoades and Dempsey, 1985). We grew homozygous Ab10-Df(K) plants and confirmed that they are striated (see below). In addition, we Illumina sequenced the genomes of plants containing Ab10-Df(K) and two other deletion chromosomes (Ab10-Df(L) and Ab10-Df(M) (Hiatt and Dawe, 2003b)) that do not have the *sr2* phenotype, and aligned the short read data to the Ab10 reference (Liu et al., 2020). The results showed that nearly all of the shared region is present in Ab10-Df(K), suggesting that *sr2* must lie at the end of the shared region of both haplotypes, presumably within a very small region or rearrangement (Figure 1, Supplementary Figure 1) (Rhoades and Dempsey, 1985)).

**Figure 1.**
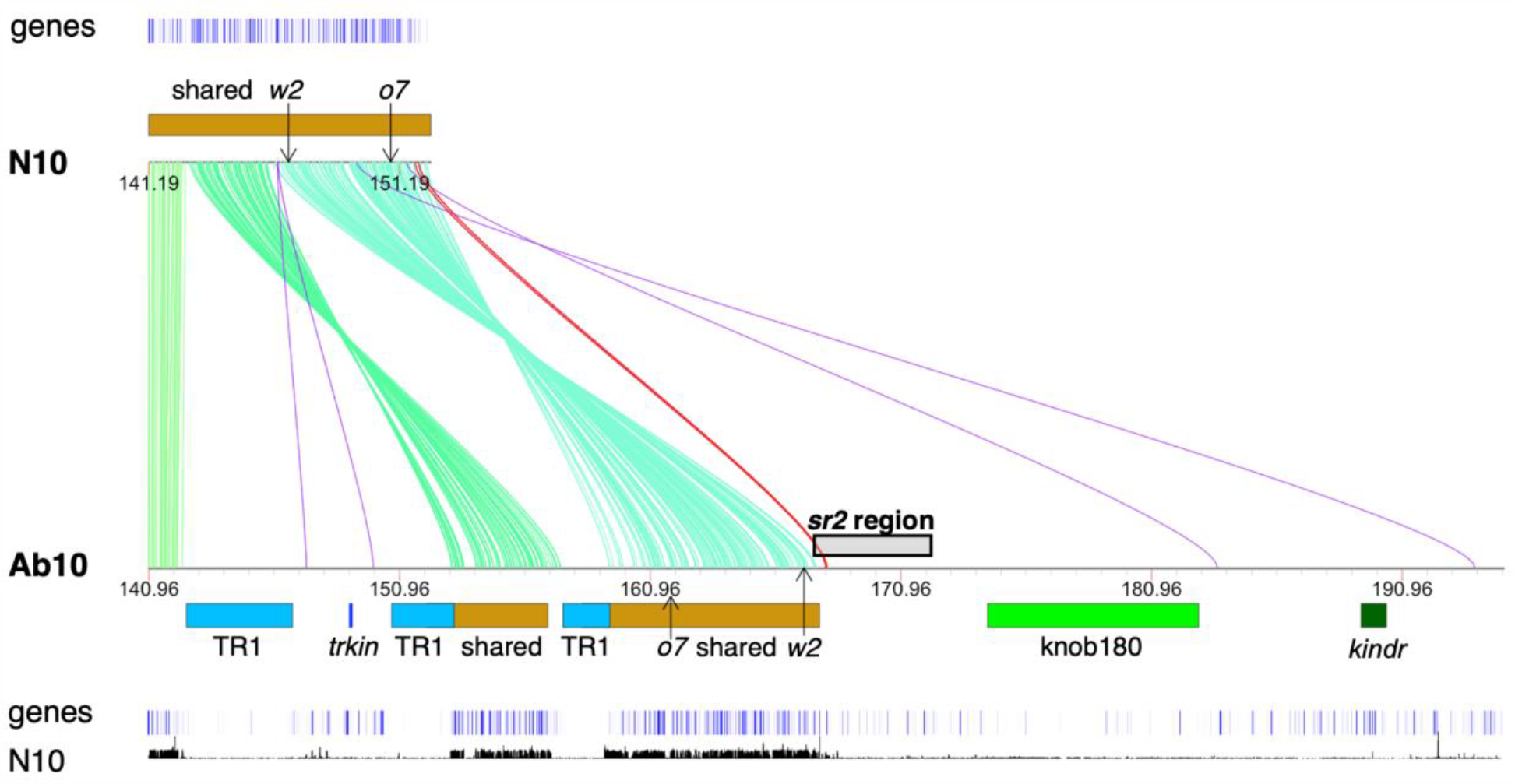
Orthologs between the N10 and Ab10 assembly. Lines indicate orthologs between N10 and Ab10 determined by OrthoFinder (Emms and Kelly, 2019). Shades of green indicate expected orthologs, red and purple indicate unexpected orthologs that could or could not be *sr2*, respectively. Location of genetic markers with known physical position, *o7* and *w2*, are shown (Wang et al., 2011; Udy et al., 2012). TR1 (light blue) and knob180 (bright green) are maize knob types. *Trkin* (dark blue) and *kindr* (dark green) are kinesin proteins responsible for the preferential transmission of Ab10 (Dawe et al., 2018; Swentowsky et al., 2020). Shared (brown) indicates the regions of known homology between Ab10 and N10. The *sr2* region is where the *sr2* gene has been mapped on Ab10, as defined by the breakpoints of Ab10-Df(K) and Ab10-Df(M) (Hiatt and Dawe, 2003b) (Supplementary Figure 1). Annotated genes are indicated as blue vertical bars (Liu et al., 2020; Hufford et al., 2021). The plot at the bottom shows short reads from B73 (which has N10) mapped to the Ab10 reference assembly with a mapping quality greater than or equal to 20 (Hufford et al., 2021). This alignment shows the traditionally defined shared region.

### Identification of a duplicated and inverted region in Ab10

We used OrthoFinder and BLAST to detect ortholog gene pairs within the *sr2* candidate region on both N10 (distal to *o7*, (Wang et al., 2011)) and Ab10 (between the Ab10-Df(K) and Ab10-Df(M) breakpoints). This approach revealed six homologous gene pairs (Figure1, Supplementary Table 1). Surprisingly, we found that this region is also present within the larger inversion proximal to the Ab10-Df(K) breakpoint on Ab10. The results suggest that a segment carrying the six genes was duplicated and inserted distal to the Ab10-Df(K) breakpoint (Figure 2, Table 1). The fact that it is a duplication helps to explain why it was not visible in alignments that display one-to-one homology relationships (Liu et al., 2020; Song et al., 2022). The OrthoFinder output and our own analysis of the structure and transcripts of each gene indicate that only 7 of the total 12 genes are functional (Table 1). In the terminal duplicated segment, there are five *sr2* candidate genes, which we will refer to in the next two sections as genes 1, 2, 4, 5, and 6. The reference names for these genes can be found in Table 1.

**Table 1.**
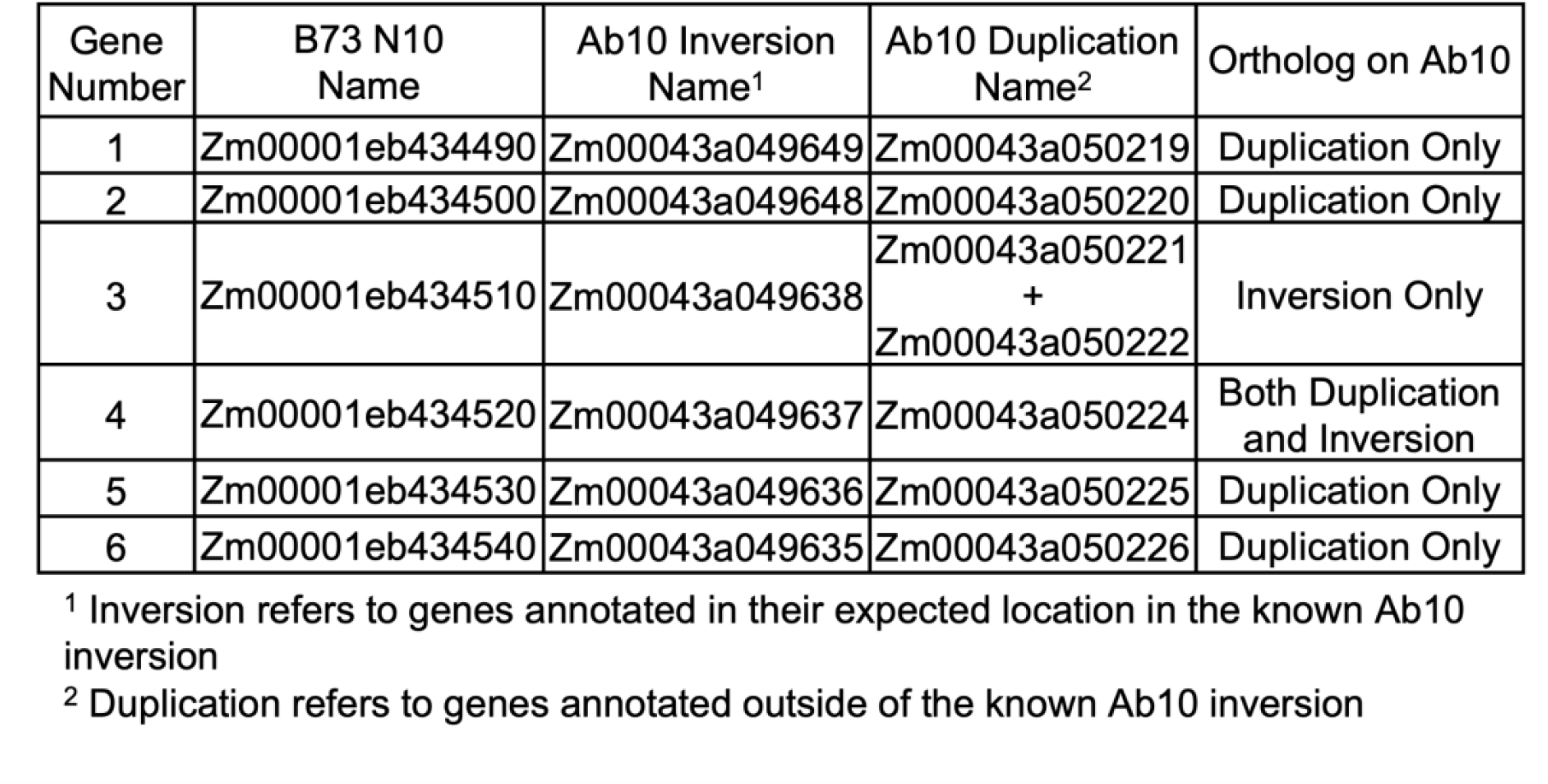
Names of genes involved in Ab10 duplication in all locations.

**Figure 2.**
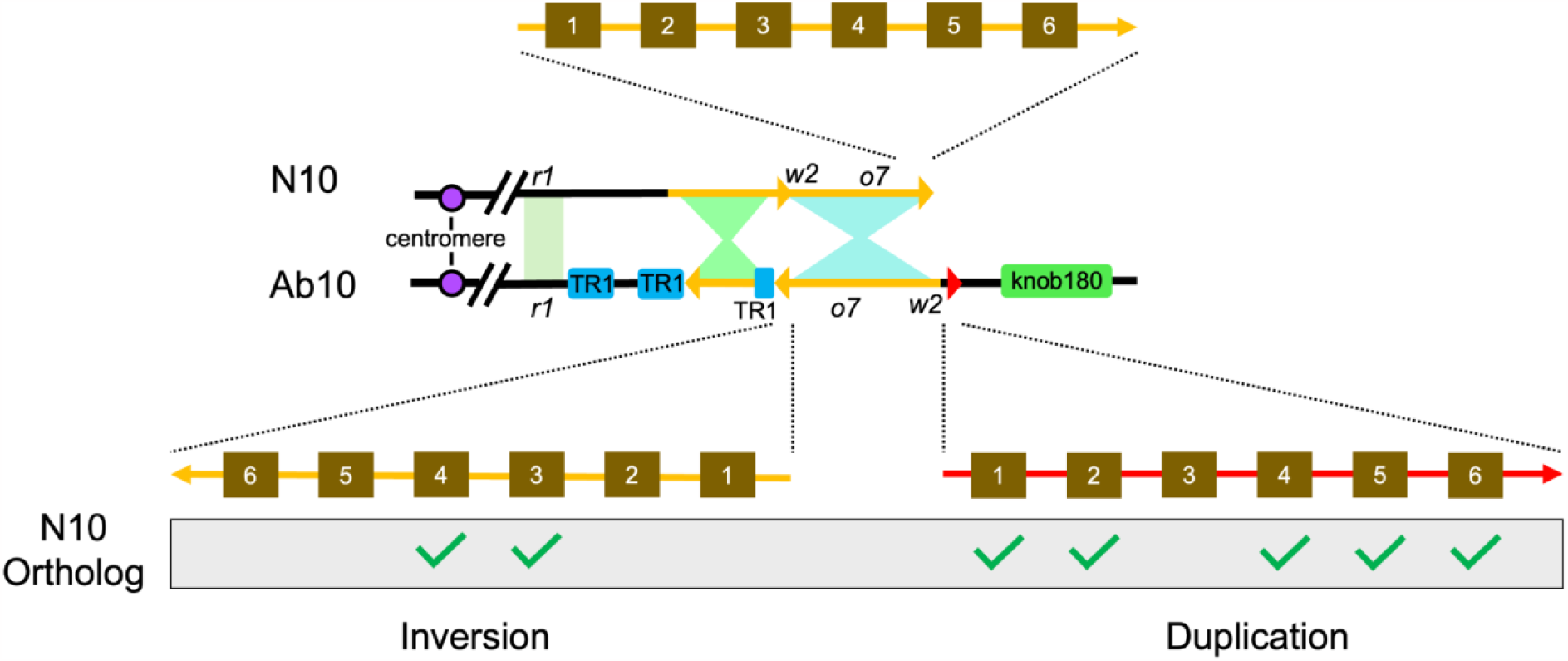
Duplicated region on Ab10. Cartoon representation of the duplicated region on Ab10 identified via OrthoFinder and BLAST (Emms and Kelly, 2019; Camacho et al., 2023). Shaded regions between N10 and Ab10 indicate regions of homology where the hourglass shape indicates an inversion. TR1 and knob180 are maize knob types and *r1* is a kernel and plant color locus marking the edge of the Ab10 haplotype. Location of genes with known physical position, *o7* and *w2*, are shown (Wang et al., 2011; Udy et al., 2012). Brown squares indicate an annotated gene (Liu et al., 2020; Hufford et al., 2021). Green checks indicate genes that appear functional.

### RNA-seq of plants homozygous for the sr2-ref reference allele implicates gene 1 as the most likely sr2 candidate

Given that the *sr2-ref* mutation has a similar phenotype to Ab10-Df(K), it seemed possible that the *sr2-ref* allele may also be associated with reduced expression. We performed a differential expression analysis using mRNA from leaf tissue. Because the genetic background of the *sr2-ref* allele is not known, we used the W22 inbred as the negative control. We aligned the *sr2-ref* and W22 RNA-seq data to the B73 v5 reference genome and performed differential expression analysis. Of the six duplicated genes, only gene 1 and gene 2 showed significant differential expression between *sr2-ref* and W22 (Figure 3). Expression of gene 2 was 44% higher in *sr2-ref* than in W22, while the expression of gene 1 was dramatically reduced to only 2% of the levels observed in the W22 inbred (Supplementary Figure 2).

**Figure 3.**
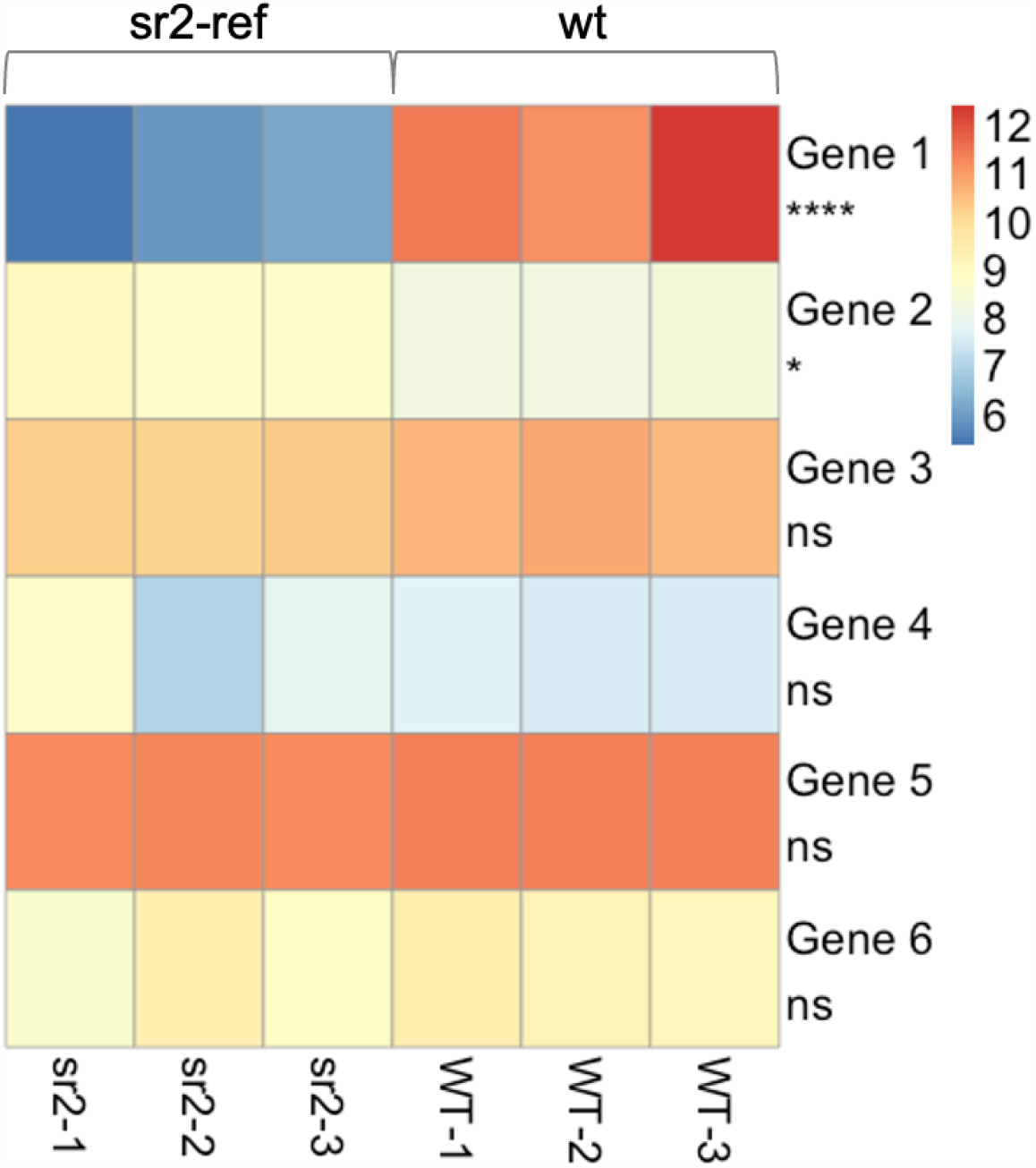
Differential expression of duplicated genes between *sr2* and wild type plants. Gene numbers refer to those defined in Table 1. Color represents log2 transformed expression value for each gene. ****= less than 0.0001, *= less than 0.05, ns = not significant.

De novo assembly of the (relatively few) gene 1 (Zm00001eb434490) transcripts in *sr2-ref* suggest that there is an insertion that often creates a chimeric transcript that omits the first exon (Supplementary Figure 3). The insertion itself appears to be a chimera of the second through fourth exons of the CASP-like protein 4A2 (Zm00001eb231550) and a DNA transposon (Zm10271_AC186904_1). The transposon Zm10271_AC186904_1 is annotated as a Robertson’s mutator (Mu) element, but we can find no homology to Mu in the terminal inverted repeat sequence (TIR). If the chimeric transcript from *sr2-ref* is translated, only a portion of the encoded protein would be homologous to the wild type gene 1 protein. There is also a very small amount of full length gene 1 transcript in *sr2-ref*, but it is >99% reduced relative to W22 (Supplementary Figure 3).

To confirm the presence of the insertion in the gene1 (Zm00001eb434490), we designed three pairs of primers based on transcript isoforms, each with a forward primer in the insertion and a reverse primer in the gene (Supplementary Table 2). All three primer pairs produced amplicons from *sr2-ref* DNA, but not from W22 control. We then Sanger sequenced the shortest amplicon (794 bp, forward primer matching the part of the insertion homologous to CASP like 4A2). Its sequence included not only CASP like 4A2 sequence, but also the 3’ 309-bp TIR of Zm10271_AC186904_1 that had been spliced out of all detected transcripts (the 5’ TIR is present in some transcripts, see Supplemental Figure 3, isoform 5). It also revealed its precise insertion point in the first intron of gene 1 (Supplementary Figure 4). These results suggest that the reduced expression in *sr2-ref* is because a DNA transposon carrying a truncated piece of CASP like 4A2 inserted into its first intron and disrupted both transcription and splicing.

### Complementation tests using transposon-induced alleles confirm that sr2 is gene 1

To further test which of the candidate genes were *sr2*, we carried out complementation tests with Mu insertions for genes 1, 2, 4, and 5 from the UniformMu collection ((Settles et al., 2007), there are no mutants for gene 6). Each of the mutant alleles contained a Robertson’s mutator element within the first exon (Supplementary Table 3). Unfortunately the genetic background of these lines is not ideal for testing recessive alleles of *sr2*. All UniformMu lines carry an allele of the *colored1* gene known as *R1-r:standard* (Settles et al., 2007; McCarty et al., 2013). *R1-r:standard* is tightly linked to a dominant allele of *inhibitor of striate1* (*Isr1)* that inhibits the *sr2* mutant phenotype (Kermicle and Axtell, 1981; Park et al., 2000). Although it is theoretically possible that *Isr1* was recombined from *R1-r:standard* during the preparation of the UniformMu lines (McCarty et al., 2005), it seems highly unlikely, as *Isr1* (Zm00001eb429350 on chr10:141210513-141213010) is only ∼20 kb from the P component of the complex *r1* locus (Zm00001eb429330 on chr10:141187279-141196584) (Walker et al., 1995; Park et al., 2000). One copy of *Isr1* reduces the striping in a homozygous *sr2* line, while two copies nearly eliminates the *sr2* phenotype. In plants homozygous for both *Isr1* and *sr2*, leaves are thinner, and white stripes are only observed at the edges of leaf margins (Park et al., 2000).

To generate material for the complementation tests we crossed lines carrying Mu alleles of genes 1, 2, 4, and 5 to *sr2-ref*. The expectation was that if one of the genes is the *sr2* gene, the Mu allele/*sr2-ref* heterozygote for that gene would show a striated phenotype. We also self-crossed the UniformMu lines to obtain plants that were homozygous for Mu alleles of each of the four genes. Phenotypic analyses of the progeny of these crosses, along with positive and negative controls, were carried out in both the greenhouse and field. Homozygous *sr2-ref* and Ab10-Df(K) had white stripes on leaf sheaths and blades in both environments, with *sr2-ref* being more striped in the field (Supplementary Figure 5).

The complementation data indicate that gene 1 (Zm00001eb434490) is the *sr2* gene. We observed leaf margin striping in the complementation tests for gene 1 and in plants homozygous for the gene 1 Mu allele (called *sr2-Mu* here forward), but not in lines carrying Mu alleles of the other three genes (Figure 4, Table 2). The *sr2-Mu* homozygous plants grew poorly and had thin leaves (Figure 4F), particularly in the field where they died before striping is normally visible. However, when they grew to maturity in the greenhouse, *sr2-Mu* homozygous plants consistently had white stripes on the edges of sheath margins (Table 2, Figure 4).

**Table 2.**
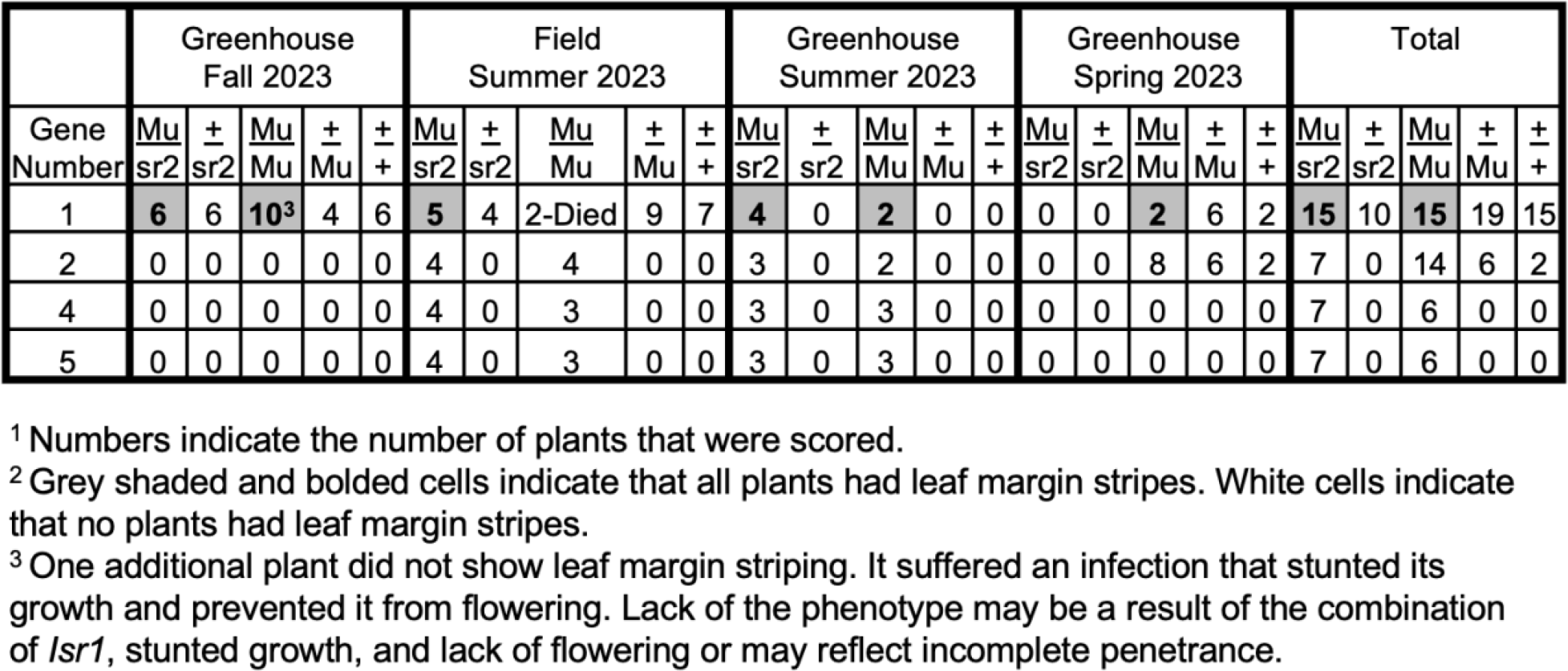
Summary of leaf margin striping phenotypes in crosses with Mu alleles^1,2^.

**Figure 4.**
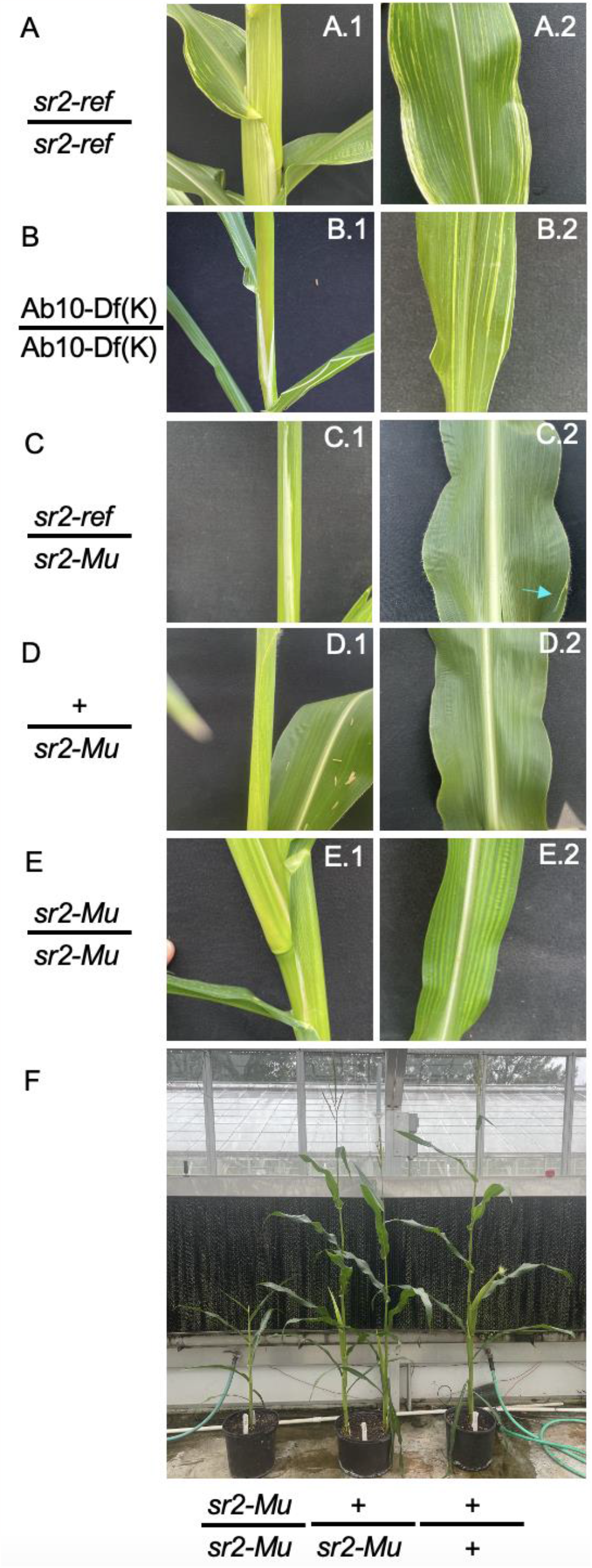
Phenotypes of *sr2* mutants. A. Homozygous *sr2-ref* plants grown in the field. B. Homozygous Ab10-Df(K) plant grown in the greenhouse. C. Plant heteroallelic for *sr2-ref/sr2-Mu* grown in the greenhouse. These plants typically had few small leaf margin stripes indicated by the blue arrow. D. Plant heterozygous for *+/sr2-ref* (where + is wild type) grown in the greenhouse. This plant is a sibling of the plant shown in C. E. Plant homozygous for *sr2-Mu* grown in the greenhouse. F. Sibling plants demonstrating the severe phenotype in *sr2-Mu*/*sr2-mu* homozygous plants relative to siblings.

The *sr2* gene is annotated as a “BTB/POZ domain protein TNFAIP protein.” However *sr2* has no homology to either the BTB/POZ domain or TNFAIP1. Rather, *sr2* is a DUF3732 domain protein, with strong (86%) protein homology to the rice *Young Seedling Stripe1* (*yss1*) gene (LOC_Os04g59570, (Zhou et al., 2017)). The fact that *sr2* and *yss1* have similar structure and function is consistent with the fact that *sr2* is a core gene in maize, found in all 26 NAM founder inbreds and is highly expressed in leaf tip tissue (Hufford et al., 2021) (Supplementary Figure 2).

## DISCUSSION

*Striate leaves2* (*sr2*) is a morphological marker that has been known since the 1940s and has played an important role in understanding the structure of abnormal chromosome 10. Comparative genomics and analysis of two independent alleles of *sr2* indicate that the causal gene is a DUF3732 domain-containing gene that is homologous to rice *yss1*. The phenotypes of *sr2* and *yss1* are similar, except that *yss1* stripes are only present in early leaves, which is not the case in *sr2*.

The rice YSS1 protein is localized to nucleoids (chloroplast genomes) and displays reduced expression of genes that are transcribed by plastid-encoded RNA polymerase (Zhou et al., 2017). In Arabidopsis, the homolog of *sr2* and *yss1* is AT4G33480. Biochemical data show that the AT4G33480 protein physically associates with psbA mRNA, which encodes a component of the Photosystem II reaction center in chloroplasts (McDermott et al., 2019). These data are consistent with the interpretation that SR2/YSS1/AT4G33480 functions at the level of chloroplast gene expression (Zhou et al., 2017). Our data further show that SR2 is unlikely to be absolutely required for chloroplast function (at least in maize), because homozygous Ab10-Df(K) plants lack the *sr2* gene yet still grow to maturity. Similar to *ij1, sr2* appears to promote faster and/or more accurate chloroplast biogenesis (Han et al., 1992).

The *sr2* phenotype is highly variable. Homozygous *sr2-ref* plants grown in the same environment can vary from having very few to very many stripes (Supplementary Figure 5). The degree of striping is also environmentally sensitive, with full sibling *sr2-ref* plants grown in the field having more severe leaf margin striping than those grown concurrently in the greenhouse (Supplementary Figure 5). The powerful effects of *Isr1* in reducing the *sr2* phenotype illustrates that there are also genetic modifiers of *sr2* (Park et al., 2000). The *Isr1* gene suppresses the growth of white tissue in *sr2* and other white striped mutants such as *ij1*. Our observation that plants homozygous for *sr2-Mu* tend to have thin leaves and few stripes can be at least partially explained by the presence of *Isr1* in the UniformMu background (Figure 4E, F). The heavy reliance on the UniformMu resource in recent years may have inadvertently limited the identification of the multiple other *striated leaves, iojap striping* and *japonica striping* loci that have yet to be described at the molecular level (Gerald Neuffer et al., 1997).

## METHODS

### Plant material and growth

Ab10-Df(K) was obtained from Marcus Rhoades and backcrossed to the W23 inbred (Hiatt and Dawe, 2003a). Ab10-Df(L) and Ab10-Df(M) were identified in our lab (Hiatt and Dawe, 2003b). Seeds carrying *sr2-ref* (X16D) and all Mu alleles were obtained from the Maize Genetics Cooperation Stock Center in Urbana, IL (Supplementary Table 3). Most experiments were conducted in the UGA Botany greenhouses in Athens GA. We also grew plants in an outdoor field site in Athens GA in April-June 2023.

### Sequencing deletion lines

We collected young leaf tissue from single plants homozygous for Ab10-Df(K), Ab10-Df(L), and Ab10-Df(M) and extracted DNA with the IBI Genomic DNA Mini Kit (Plant) (IB47230) using the GP1 buffer and 16,000xg for all centrifuge steps. We used a Kapa HyperPrep Kit to prepare the sequencing library (KK8580) and adapters from Netflex DNA Barcodes (Nova-520996). Sequencing was performed by GENEWIZ from Azenta using a HiSeq 4000. We trimmed the resulting reads using cutadapt version 2.7 (Martin, 2011), mapped the reads to the Zm-B73_AB10-REFERENCE-NAM-1.0 reference (Liu et al., 2020) using BWAmem 0.7.17 (Li and Durbin, 2009), and removed duplicates using picard version 2.16.0 (Tools, 2018). We determined the breakpoints by filtering the primary alignments with a MAPQ>=20, counting the number of reads over each bp using IGV tools, and visually inspecting the resulting files (Robinson et al., 2011). N10 reads shown in Figure 1 and Supplementary Figure 1 are B73 ∼30X Illumina short reads from the NAM project (Hufford et al., 2021). Plots were made using R v4.3.1.

### Identifying and analyzing orthologs in the duplicated region of Ab10

We selected the protein sequence for the longest isoforms of the annotated genes between the *colored1* (*r1*) gene (Zm00001eb429330, B73 v5 annotation) and the ends of the long arm of chromosome 10 in the the Zm-B73-REFERENCE-NAM-5.0 (https://www.maizegdb.org/genome/assembly/Zm-B73-REFERENCE-NAM-5.0) and Zm-B73_AB10-REFERENCE-NAM-1.0 (https://download.maizegdb.org/Zm-B73_AB10-REFERENCE-NAM-1.0/) assemblies (Liu et al., 2020; Hufford et al., 2021). These files were used to run orthofinder v2.5.2 using default parameters (Emms and Kelly, 2019). Orthofinder identified five genes that were duplicated at the end of the shared region. BLAST v 2.2.26 (Camacho et al., 2023) was used to identify a sixth gene within the duplication. Plots were made using R v4.3.1.

We analyzed the transcripts for all six genes in the Ab10 inverted, Ab10 duplicated and N10 regions using the multiple sequence aligner and Clustal Omega in Geneious Prime v2022.0.2 (https://www.geneious.com/). We used the coding sequence (CDS) data published with the B73 and B73-Ab10 reference genomes (Liu et al., 2020; Hufford et al., 2021). For gene 1 within the inversion, all transcript isoforms were truncated due to an unknown insertion in exon 4 which caused a frameshift and premature stop codon in exon 5. For gene 2 within the inversion, we found that all transcript isoforms either lacked significant homology to the N10 gene or were truncated for unknown reasons. For gene 3, we used Clustal Omega to determine that the duplication homologs are missing 288bp of coding sequence and are unlikely to be functional. For gene 4, we used Clustal Omega to determine that both the Ab10 inverted and duplicated copies are similar to the N10 homolog and are likely both functional. For gene 5, we used Clustal Omega to determine that the Ab10 inversion homolog has one isoform that is severely truncated for an unknown reason and one that results in the insertion of a proline within the PaaI thioesterase domain (Marchler-Bauer et al., 2017). This may or may not disrupt function. For gene 6, we used Clustal Omega to determine that both Ab10 homologs are longer than the N10 counterpart and that both the Ab10 inversion and duplication homologs have a similar amount of homology to the N10 copy. This may or may not disrupt function in the Ab10 homologs. These data in conjunction with the results from OrthoFinder indicate that genes 1, 2, 4, 5, and 6 are candidates for *sr2* (Table 1).

### RNA-seq of plants homozygous for sr2-ref

#### RNA isolation and Sequencing

We collected mature leaf tissue from homozygous *sr2-ref* and W22 individuals grown side by side in the UGA Botany greenhouse and immediately froze it in liquid nitrogen. RNA was extracted using the IBI Total RNA Mini Kit (Plant) (IB47340) and cDNA was prepared using BioRad iSCRIPT Transcription Supermix for RT-PCR (1708891). GENEWIZ from Azenta performed a library preparation with PolyA selection using an NEBNext Ultra II RNA Library Prep Kit followed by paired end 150bp illumina sequencing on a NovaSeq 6000.

#### Differential Expression Analysis

We trimmed reads using Trimmomatic version 0.39, and checked quality before and after using FastQC version 0.11.9 (Bioinformatics, 2014; Bolger et al., 2014). We aligned reads to the Zm-B73-REFERENCE-NAM-5.0 reference using HISAT2 version 2.1.0 using default parameters (Kim et al., 2019; Hufford et al., 2021). Using HTSeq version 0.13.5 with default parameters, we determined the number of reads mapped to each annotated feature (Anders et al., 2015). All genes with fewer than 10 reads were removed. We used DESEQ2 version 1.38.3 to perform a differential expression analysis with default parameters. Plots were made using R v4.3.1.

#### *De novo assembly of* sr2-ref *transcripts*

We performed a Trinity v2.10.0 de novo transcriptome assembly on pooled data for all three biological replicates of *sr2-ref* (Haas et al., 2013). BLASTv 2.2.26 was used to identify isoforms with homology to the *sr2* gene. Some of the isoforms also contained sequences with homology to CASP-like protein 4A2 (B73 v5 annotation Zm00001eb231550) and L-aspartate oxidase (B73 v5 annotation Zm00001eb231540). The relative abundance of each isoform was determined using Kallisto within Trinity v2.8.4 (Haas et al., 2013; Bray et al., 2016). All transcript assemblies were then aligned to the B73 *sr2* reference gene (Zm00001eb434490) using Geneious Prime v2022.0.2 (https://www.geneious.com/) MiniMap2 with default parameters followed by minimal manual curation.

The *sr2-ref* allele insertion was confirmed via PCR using primers to CASP like 4A2 and *sr2* (Supplementary Table 2). The shortest PCR product was Sanger sequenced by Eton Biosciences .

### Genotyping

All genotyping DNA extractions were performed using a CTAB protocol (Clarke, 2009). Polymerase chain reactions were performed using Promega GoTaq Green Master Mix (M7123). The wild type locus for each gene was detected using gene specific primers (Supplementary Table 2). The Mu allele for each gene was detected using a primer to the Mu terminal inverted repeat (5’-GCCTCYATTTCGTCGAATCCS-3’) and either a forward or reverse gene specific primer. All genotyping reactions used the following temperature profile: Hold 95°C, 2min 95°C, 30(30sec 95°C, 30sec 60°C, 45sec 72°C), 5 min 72°C.

### Complementation tests using Mu-induced alleles

For genes 2, 4, and 5 we produced seed that was homozygous for the Mu insertion of interest and crossed these to *sr2-ref* to produce seed that was all heterozygous for the insertion and *sr2-ref*. For gene 1, we generated segregating populations of mutant and wild type by self crossing a plant heterozygous for the Mu insertion and crossing this plant to the *sr2-ref* tester.

Plants were grown in both the greenhouse and field as detailed in Table 2. In the field we randomized plant location and surrounded experimental plants with buffer corn to limit any environmental effects. In all cases we grew *sr2-ref* and Ab10-Df(K) plants alongside the Mu bearing plants to confirm that the conditions were appropriate to see the phenotype.

## ACCESSION NUMBERS

All code is available at https://github.com/dawelab/Striate_Leaves2. Sequencing data from this project has been deposited to the SRA under bioprojects PRJNA1019348 and PRJNA339461. Relevant gene numbers can be found in Table 1, and Supplementary Figure 3.

## ACKNOWLEDGEMENTS

We thank Yibing Zeng for help with accessing expression data for *sr2* in the NAM lines and the Maize Genetics Cooperation Stock Center for maize stocks. This work was supported by a NIH training grant (T32GM007103) and NSF fellowship to MJB (2236869) as well as an NSF grant to RKD (1925546) and technical expertise from the Georgia Advanced Computing Resource Center.

## AUTHOR CONTRIBUTIONS

MJB and RKD designed the research. MJB, JIG, and MC performed research; MJB analyzed data. MJB and RKD wrote the paper.

