## Supplemental Tables and Figures for "The maize *striate leaves2* (*sr2*) gene encodes a conserved DUF3732 domain homologous to the rice *yss1* gene"

**Supplementary Table 1. Orthologs outside of regions of known Homology between Ab10 and N10.**

| B73 Gene Name | Ab10 Gene Name | Gene Number |
|---|---|---|
| Zm00001eb431630 | Zm00043a049151 | NA[1] |
| Zm00001eb431660 | Zm00043a049258 | NA |
| Zm00001eb433140 | Zm00043a051035 | NA |
| Zm00001eb434390 | Zm00043a050572 | NA |
| Zm00001eb434490 | Zm00043a050219 | Gene 1 |
| Zm00001eb434500 | Zm00043a050220 | Gene 2 |
| Zm00001eb434520 | Zm00043a050224 | Gene 4 |
| Zm00001eb434530 | Zm00043a050225 | Gene 5 |
| Zm00001eb434540 | Zm00043a050226 | Gene 6 |

[1] Genes labeled NA were not considered *sr2* candidates because they are located outside of the mapped *sr2* region (the purple lines in Figure 1).

**Supplementary Table 2. Gene specific primers[1].**

| Name | F sequence | Target | Name | R sequence | Target |
|---|---|---|---|---|---|
| Sr_490_F | GTCGAGATAGAAGGGGCGAG | Zm00001eb434490 | Sr_490_R | GCGAGCTCCGGAATACTCTC | Zm00001eb434490 |
| Sr_500_F | CAACATGTGGAGCGACCAAC | Zm00001eb434500 | Sr_500_R | GGAACCCGATCTAAGAGGCG | Zm00001eb434500 |
| Sr_520_F | CAGGAAGAGCACACGGAAGA | Zm00001eb434520 | Sr_520_R | CAGACAAGGCGGATTGACCT | Zm00001eb434520 |
| Sr_530_F | GTAGTCTGGAGGGCACGTTG | Zm00001eb434530 | Sr_530_R | GAAAGCGTAAAGACGTGGCG | Zm00001eb434530 |
| JIG-419 | TAGGCTTCCTCCTGCTCTC | Zm00001eb231550 4th exon | JIG-420 | TGCTTGAGTTGGTCGTCATC | Zm00001eb434490 4th exon |
| JIG-421 | CTAACGGCTGCTGGTCTG | unique to sr2-ref transcript | JIG-422 | GCTACCCAGCCACTGAAAT | Zm00001eb434490 3rd exon |
| JIG-423 | TGCTGGTCTGCGAGGAT | unique to sr2-ref transcript | JIG-424 | GCCACTGAAATTTGTGCTCTTG | Zm00001eb434490 3rd exon |

[1] All primers are shown in the 5' to 3' orientation.

**Supplementary Table 3. Mu alleles used in this study.**

| Gene Number | B73 Gene Name | Uniform Mu Stock | Mu of interest | Location |
|---|---|---|---|---|
| 1 | Zm00001eb434490 | UFMu-08182 | mu1058934 | 1st exon |
| 2 | Zm00001eb434500 | UFMu-12931 | mu1089572 | 1st exon |
| 4 | Zm00001eb434520 | UFMu-03495 | mu1037653 | 1st exon |
| 5 | Zm00001eb434530 | UFMu-03745 | mu1037651 | 1st exon |

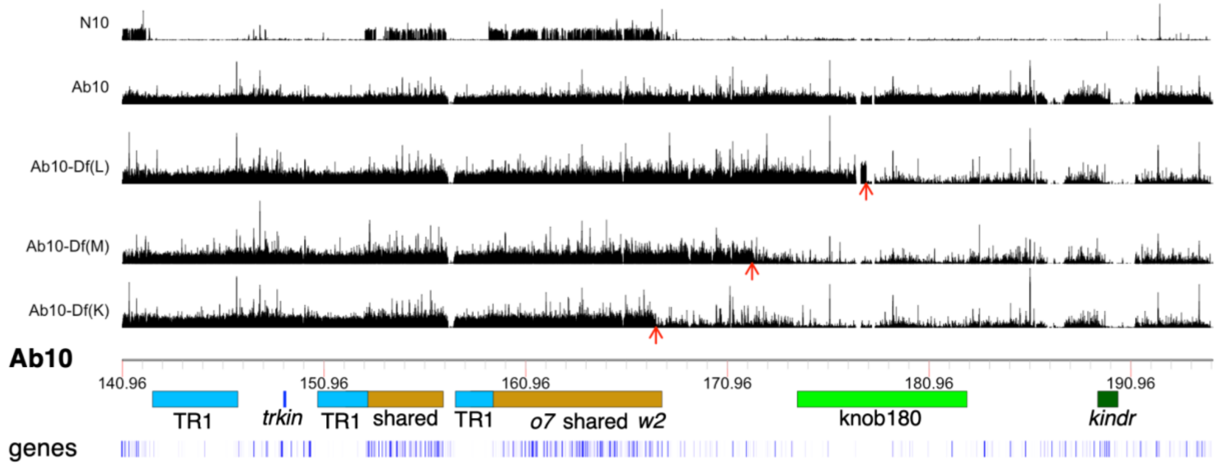


**Supplementary Figure 1. Ab10 deletion breakpoints.** Black bar plots show average read count calculated in 1 kb windows for Illumina paired end reads mapped to the Ab10 reference using BWAmem filtered to a MAPQ of 20 and only primary alignments. Red arrows indicate the presumed breakpoint. Location of genes with known physical position, *o7* and *w2*, are shown (Wang et al., 2011; Udy et al., 2012). TR1 and knob180 are maize knob types. *trkin* and *kindr* are kinesin proteins responsible for the preferential transmission of Ab10 (Dawe et al., 2018; Swentowsky et al., 2020). Shared indicates regions of known homology between Ab10 and N10. Blue bars indicate annotated genes (Liu et al., 2020).

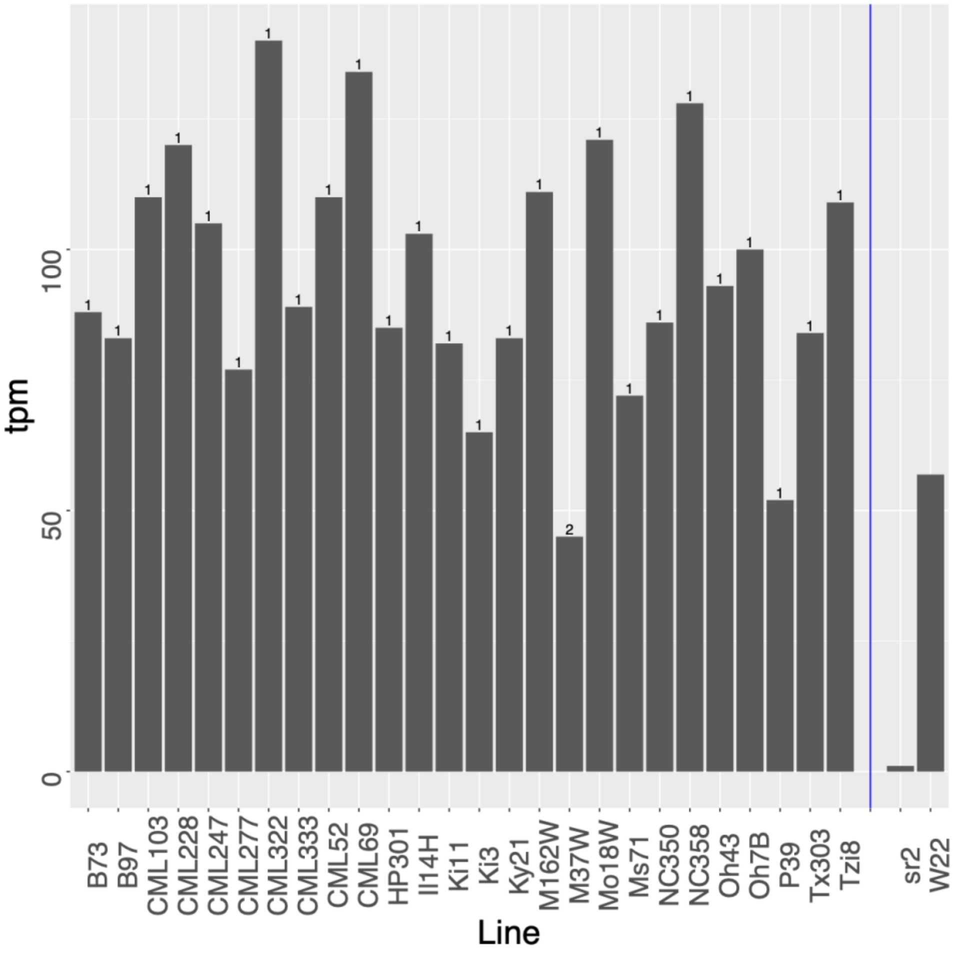


**Supplementary Figure 2. Expression of *sr2*.** Leaf tip expression values (bar height) and copy number (number above bar) for *sr2* in all NAM founders (Hufford et al., 2021). Data to the right of the blue line are leaf tip expression data from this project, copy number is unknown for these lines.

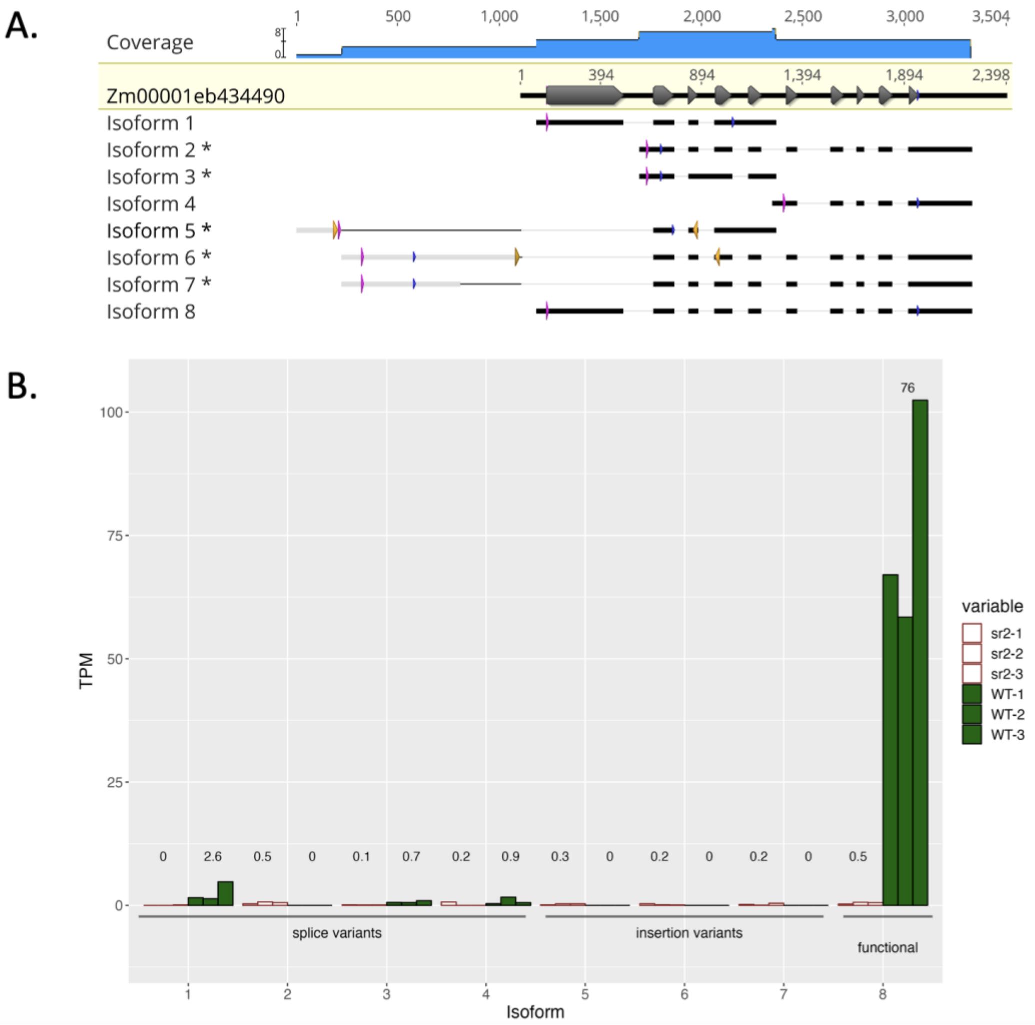


**Supplementary Figure 3. The *sr2-ref* allele contains an insertion.** A. The first line shows the gene model from B73, where grey arrows indicate exons. Isoforms 1 through 8 are derived from a Trinity de novo transcriptome assembly of *sr2-ref* (Haas et al., 2013). * indicates there is a frame shift relative to the wild type in that isoform. Pink segments indicate the first ATG, blue segments indicate the first in-frame stop codon, gold segments indicate primers. Light grey boxes indicate sequence that is different from the *sr2* B73 reference, black indicates strong homology to *sr2*. Isoform 5 includes sequence with homology to the DNA transposon Zm10271_AC186904_1. Isoforms 6 and 7 include sequence with homology to CASP-like protein 4A2 (Zm00001eb231550). None of the isoforms include the 3’ TIR of Zm10271_AC186904_1, indicating it is reproducibly spliced out of the transcript (Supplemental Figure 4). B. Transcripts per million (TPM) for each isoform in each biological replicate of *sr2-ref* and wild type leaf tissue. The number above each set of three bars represents the average.

**JIG-419**

**TAGGCTTTCCCTCTGCTCTCCTCCAG**GTACGGGAAGTCCTCGGTGAAGTGCAGGCCGCGGCT

CTCGCGCTTGGCCAGCGCGCTCCTGACGACGTGCAGACGAGCGCCGCCGCTCCTGCTCGGGC

GCCGGCTCGGCACCTGCTCGGGCCGCCGGCTCGCTCGTCAGGGAGAGAGCGCTGGGGGAGAG

GAGCGCCCGTGGATCCGTCGTAAGAAACGGACTGCCGTAACACCTAGGTTAAATCGAGCTAA

ATCTGACTCAGATACTACAATTCCAGAGTTCAACGGATTATAATACTAAAATTCCAAATGTG

ATGTTGGGAATACCTATTTTCAGAAGCCCAGTGCTGACGATATCACTGTTCCAAAAAGCTCC

ATCCCGAGCTTTTTGGAACCGTGATATCCCCAGCTTAATATTTTCTTTCTTTCTGTCGTCAC

AAATGAAATCAAGCCAAACCAACATTGTGATATGCAG**ATCGACGACACCAACAACCGCCCCT**

**TGGAGTGCATAATCAGGAGAGTATTCCGGAGCTCGCAGGACCACGACTGCATGCTTCTCTGC**

**CCAGTTGACAT**GTAAGTGACACTCTCTCTTTTTTGTACTCATACTCATCGCAGACACCCAAG

TCACACTCTCTCTCTCTTACA**GGCCTGTGCAGGTCCTCAAGAGCACAAATTTCAGTGGCTGG**

**GTAGCT**GTATGACTTGGCCCTCATATACATTCCTTCAACTGATTATTTCATTTATTTCCCAT

GGTCCTAATA*CCTGTTTATTTATTTAGGTCGATGACGACCAACTCAAGCA*

**JIG-420**

**Supplementary Figure 4. The *sr2-ref* insertion is in the first exon.** Sanger sequenced amplicon of JIG-419 and JIG-420 (Supplementary Table 4) from *sr2-ref* DNA. Blue solid underlined text indicates homology to CASP-like protein 4A2 (Zm00001eb231550). Green dotted underlined text indicates homology to the DNA transposon Zm10271_AC186904_1. Orange not underlined text indicates homology to *sr2* (Zm00001eb434490). Bold text indicates sequence in *sr2-ref* isoform 6 (Supplementary Figure 3). Italicized text indicates sequence that was expected in the amplicon, but did not appear in the Sanger sequence. Grey highlighted text indicates primers.

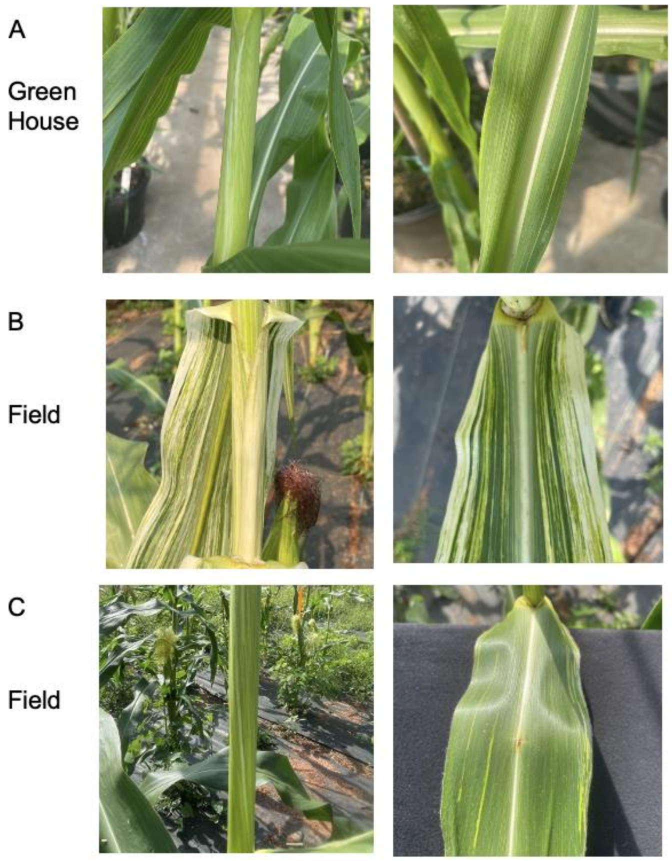


**Supplementary Figure 5. The *sr2-ref* phenotype varies with environment and between individuals.** All individuals shown are *sr2-ref* homozygous full siblings grown at the same time in Athens GA, USA. A. Plant grown in the greenhouse with weak leaf striping. This phenotype is representative of three individuals. B. Plant grown in the field with strong leaf striping. C. Plant grown in the field with moderate leaf striping. B and C, two additional individuals grown in the field that had an intermediate leaf striping phenotype.
